# Visual perception and workload of office workers in various acoustic conditions

**DOI:** 10.1101/2023.02.15.528721

**Authors:** Joanna Kamińska, Jan Radosz, Łukasz Kapica

**Affiliations:** Department of Ergonomics, Central Institute for Labour Protection - National Research Institute, Warsaw, Poland; Department of Vibroacoustic Hazards, Central Institute for Labour Protection - National Research Institute, Warsaw, Poland

**Author notes:** Corresponding author, (JK).

## Abstract

Noise in the office work environment can negatively affect workers’ cognitive performance, number of errors made and comfort. The aim of the study was to determine the effects of various types of acoustic conditions in the mental work environment on visual perception (eye-tracking parameters) and workload. Method: In the experiment a group of 39 people aged 20 to 34 was asked to perform two eye-tracking tests (Perceptiveness and Speed Tests (PTs) and to read the text of a fictional biography, and then to answer questions about the reading). Mental workload was evaluated in each condition using NASA TLX questionnaire. The tests were performed in various acoustic conditions: variant W1 – no presentation of acoustic stimuli, variant W2 – sounds of office equipment, variant W3 – sounds of office equipment with quiet conversation in native language, variant W4 – sounds of office equipment with loud conversation in native language, variant W5 – filtered pink noise. In variants from W2 to W5 the equivalent sound level A was 55 dB. Results: The assessment of work efficiency in the reading test indicates the existence of statistically significant differences. The most errors were made during mental work with the audible sounds of office equipment with a loud conversation (Variant W4) and during mental work performed with audible filtered pink noise (W5). While reading the text, different acoustic conditions do not differentiate statistically significantly visual perception described by eye-tracking parameters. In turn, in the PTs test, statistically significant differences between the variants were found in the Digit test (average blink duration) and the Fraction test (average blink duration, average fixation duration and saccades frequency parameters). In conclusion, visual perception depends on the type of noise. Acoustic factors aggect workers’ cognitive functions, mostly in more difficult tasks.

## Introduction

The term office work is usually understood as mental work of varying complexity (from routine work that does not require much intellectual involvement to creative work with a high degree of mental engagement). These jobs require memorising information, focusing attention and performing activities at various levels of difficulty. Background noise is the most common complaint reported by employees, especially in open-plan offices. This is not surprising, considering that the concept of planning open space should be based on a compromise between the need to ensure conditions for good communication among staff and maintaining the privacy of individual employees. Noise during office work has been analysed for a long time. Employees indicated nuisance related to telephone ringing, disturbed by conversations, and loud air conditioning and office equipment [1]. Assessing the impact of noise on cognitive abilities can be valuable in reducing human error and increasing productivity in noisy workplaces such as banks or contact centers.

Many studies have confirmed the existence of a direct relationship between increased noise levels and reduced cognitive performance [2–4]. Cognitive impairment can lead to decreased performance, errors and ultimately to an increase in accidents in cognitively related occupations [5]. On the other hand, according to Keighley, acceptability was not related to the level of background noise, but was strongly inversely correlated with peak sound levels above the background noise level, suggesting that distinctive or distinct sounds were the least acceptable [6]. Other research confirmed that noise-induced cognitive impairment depends on the type of noise, sound pressure level, noise spectrum and time of exposure to noise, and the difficulty of the task [7–9].

Cognitive functions include such processes as attention, perception, memory, decision making, problem solving and reaction time [10]. Noise-induced cognitive impairments are usually thought to be distractions of attention and working memory. However, perceptual processes are the first stage that allows the collection of information and, as a result, it is possible to process and perform tasks [11]. Working memory processes refer to a limited-capacity system that provides temporary storage and handling of information needed for complex cognitive tasks such as comprehension and learning [2,12]. Research also indicate an increase in the number of errors and an increase in reaction time when performing tasks with a higher level of difficulty [13]. When the tasks are simple (1-Back), then their performance is affected by the noise level (more unfavorable acoustic conditions), while when the tasks are medium or difficult (2 and 3-Back), their performance is more strongly influenced by the type of noise, not its level. Another study also showed that noise had an impact on the deterioration of attention and short-term memory [13]. Perception is a multisensory process, and previous work makes multisensory interactions not only with object-related stimuli, but also with simple and seemingly unrelated inputs from different senses [14]. Basic neural mechanisms were examined e.g. using electrical neuroimaging (EEG), acoustic noise was found to affect the occipital alpha power and beta band coupling of the occipital and temporal sites. Task-irrelevant and continuous sounds had an amplitude-dependent effect on cortical mechanisms involved in shaping visual cortical excitability. Presented results suggest that both task related sounds and background noises not relevant to the task can induce visual enhancement [14]. Many studies have demonstrated perceptual suppression effects in single visual and auditory stimuli [15–17]. A neurophysiological study [18] showed a visual perceptual masking effect that occurred due to direct interactions of neural responses. When the perception of the target stimulus was suppressed by another masking stimulus presented immediately before or after the target, neural responses associated with the onset or shift of the target were also suppressed [19]. The conducted neurophysiological studies concerned both the facilitating and suppressing aspects of the interaction. For example, auditory stimuli have been found to amplify the perceived intensity of visual stimuli [17]. Other human brain imaging studies have also shown that auditory or visual stimuli deactivated part of the visual or auditory cortex, respectively [18]. In the study [19], it was shown that the sounds emitted by headphones deteriorated the ability to distinguish visual orientation.

Scientific research reveals the possibility to use eye tracking records to analyse cognitive processes. Eye activity was examined by analysing e.g. pupil diameter and eye blink rate or blink frequency as a cognitive function indicators [20, 21, 22]. Studies investigating the potential of using eye movements for assessing the mental workload also indicate that pupil diameter and fixation time show an increase if the mental workload increases while saccade distance and saccade speed do not show any differences [23, 24]. Other scientific literature shows that saccade distance and saccade speed can be used as an indicator of mental workload [25]. Studies [26] showed that mental load exerted by performing tasks at an imposed and increasing rate (time pressure) affects the eye movement characteristics in the subjects. Statistically significant changes were observed for the blink frequency, fixation duration and saccade frequency. The changes do not clearly suggest a change towards greater load or fatigue of the organ of vision. The subjects compensate for the limitation related to perception of stimuli and possibility to respond to them quickly with an increasing number of errors (omissions and incorrectly clicked items) and a lower number of correctly performed actions (correctly clicked items).

The aim of the research was to examine the impact of the type of acoustic conditions, typical of office work, on visual perception (eye-tracking parameters), in comparison to subjectively assessed workload (NASA-TLX questionnaire) and work efficiency.

## Methodology

### Study population

The study population included 39 payed volunteers (18 women and 21 men), mean age (SD) – 24.0(4.53) years. The inclusion criteria for participants selection were following: aged 20 to 35 years; healthy subjects with good eyesight (vision defect not greater than ± 2 dioptres); employed as mental worker. Individuals who revealed any major head injuries with loss of consciousness or concussion in the last 5 years or mental diseases or were addicted to psychotropic substances were excluded.

The study was approved by Bioethics Commission of Cardinal Stefan Wyszyński University in Warsaw, approval number: KEiB—33/2021. Prior to the commencement of the study, the participants were informed about the purpose and course of the study, and signed a consent for participation in it. As stated in the consent form, the data collected by the study are confidential and participants have the right to terminate the experiment at any time and to withdraw their provided data at any moment even after the data collection.

The participants were required to complete specific cognitive tasks under different experimental noise stimuli, and to evaluate visual perception and workload caused by the experimental noise stimuli. The effects of noise were assessed by eye-tracking recordings (visual perception), by the questionnaire NASA-TLX (workload), subjective acoustic condition assessment and work efficiency results.

### Study methods

An eye tracker from SensoMotoric Instruments (SMI) was used to record the eye movement parameters. It was integrated with a 22” screen where the tasks to be performed by the study participants were displayed. IR cameras, with an automatic system for recording the eye movement, corneal reflection and modifications in the position of the study participant’s head were used as recording equipment (Fig. 1). The tests were conducted at the sampling frequency of 250 Hz, spatial resolution 0.03°, distance between the eyes and the monitor 60 - 80 cm, and the system latency: <10 ms.

**Fig. 1.**
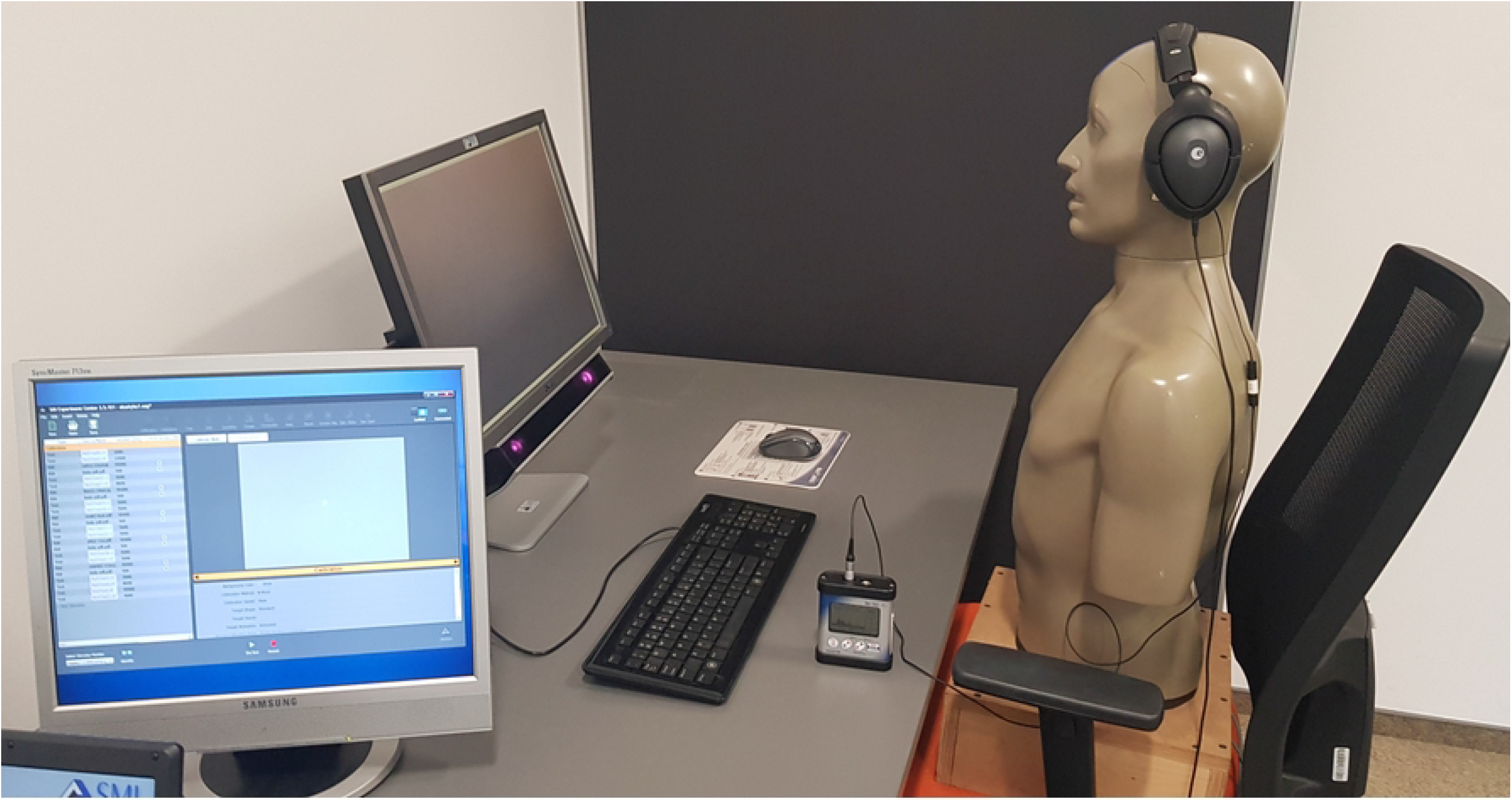
Stand for registrations.

The NASA-TLX questionnaire [27] was used for the assessment subjective, multidimensional workload. The questionnaire consists of six subscales describing the following aspects of workload: mental workload, physical workload, time pressure, productivity, effort and frustration. A question on acoustic conditions has been added to the questionnaire: How do you rate the acoustic conditions during the study?

After each variant, the participants had to assess the nuisance of the acoustic conditions - they answered the question: “How do you assess the acoustic conditions during the test?”. The subjects’ subjective feelings were rated on a scale from 0 (comfortable) to 100 (very disturbing).

### Study procedure

The task of the participants was: a) to do Perceptiveness and Speed Tests (PTs) b) to read the text of a fictional biography, and then to answer questions about the reading.

Perceptiveness and Speed Tests (PTs) consists of several sub-tests which require perceptiveness, fast working and focusing attention. Each of the sub-tests (Sub-test of Letters, Alfa Sub-test, Sub-test of Digits and Sub-test of Fractions) is based on a similar scheme and is composed of three parts (Fig. 2):

a. Screen with information and instructions for the study participant. The study participant is informed that their task is to find and count the number of occurrences of a particular character appearing among others on the screen. Depending on the sub-test, this is a Latin alphabet letter (e.g. the letter C), a Greek alphabet letter (e.g. Alpha), a digit (e.g. 3) or fraction (e.g. 2/3). The test should be performed in such a way that the characters in each line can be read from left to right, starting with the first line, and following downwards line after line. No line on the screen can be skipped and the task should be performed as quickly as possible.
b. A visual task which involves finding a set character (Latin alphabet letter, Greek alphabet letter, number or fraction) among other characters on the screen is carried out according to the instructions given in point a.
c. Screen with answers giving the number of characters found. The study participant’s task is to indicate the number of characters found (by looking at the right number) displayed on the screen.

**Fig. 2.**
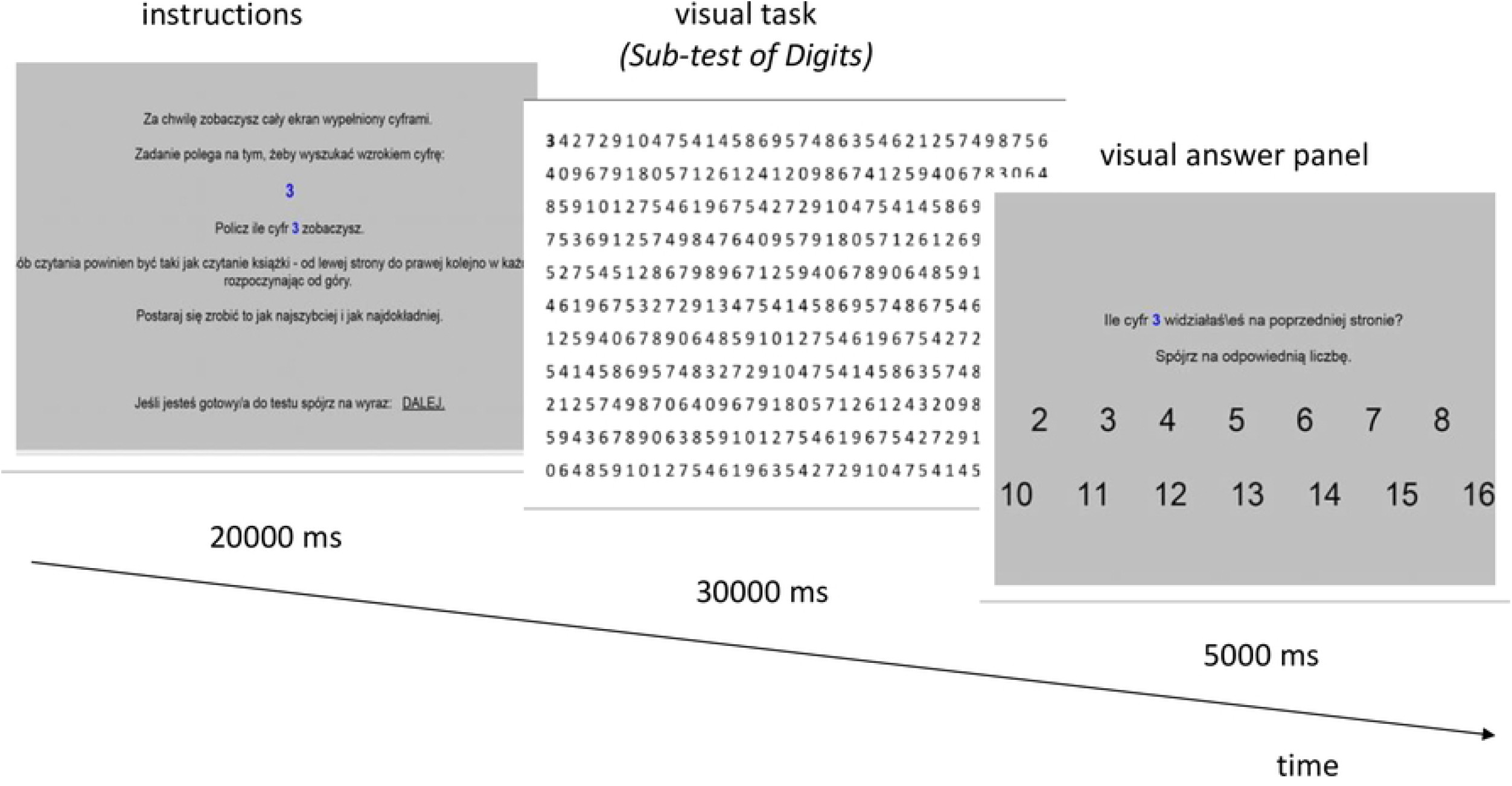
Sample screens with instructions, visual task and visual answer panel.

**Fig. 3.**
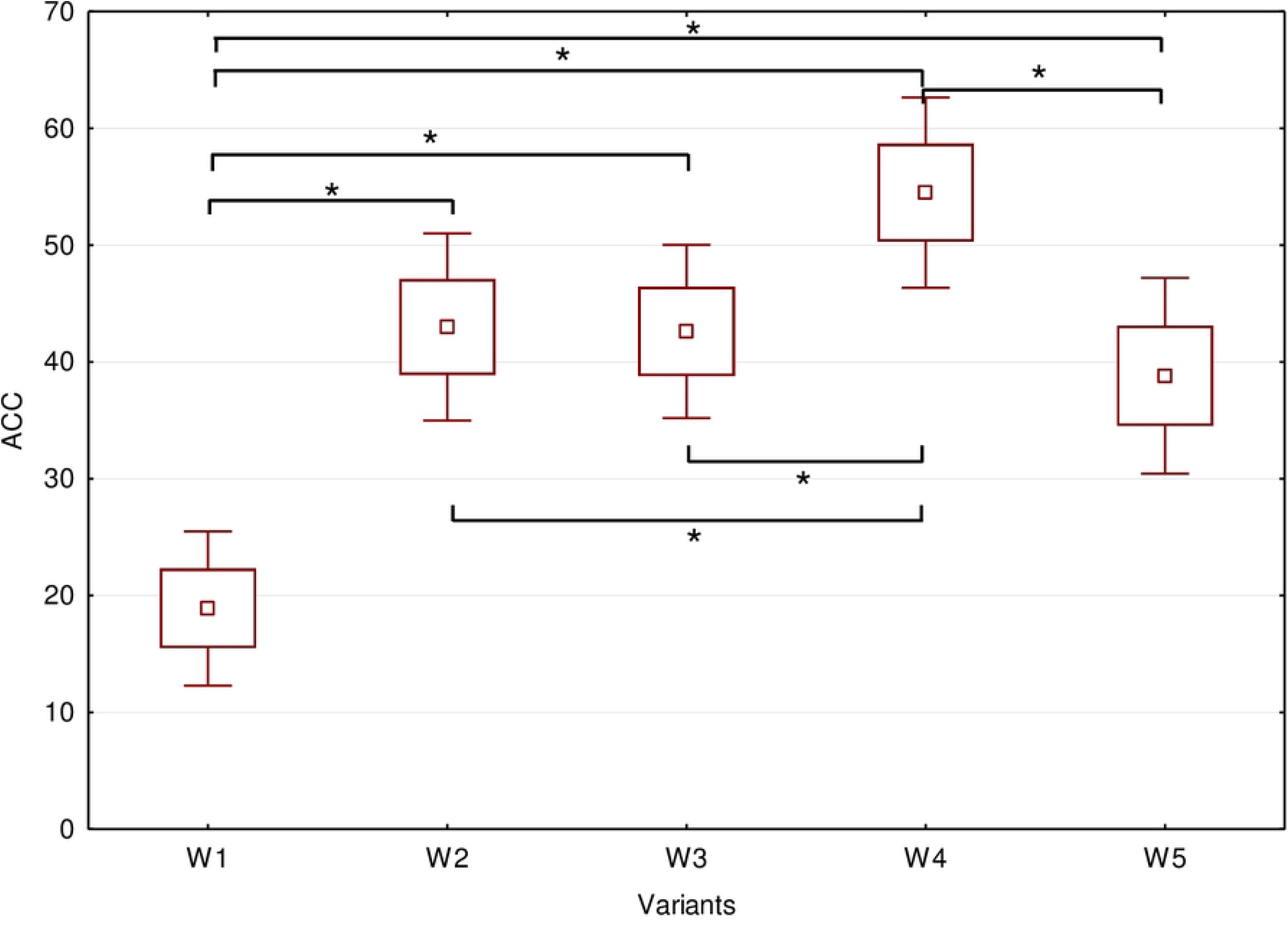
Evaluation of the subjective perception of the ACC (nuisance of the acoustic conditions)

The task ends when the study subject moves their eyesight onto the characters at the end of the last line, where a trigger is located – this is an area (invisible to the participant) which covers the last 5 characters within a paragraph. If the program registers a fixation time in this area longer than 200 ms, it moves to the next screen automatically (next sub-test or end of experiment).

Depending on the number of calibration repetitions and the time spent on each task, the total duration of the Perceptiveness and Speed Tests (PTs) is about 5–6 minutes. Prior to the test, the study participants completed a short training to eliminate the learning effect.

More severe vision defects may result in loss of some data and that is why prior to statistical analysis of the parameters, the numerical data were subject to initial processing in order to remove artefacts or such test results for which two assumptions were not met: Tracking Ratio <90% and the value of the coefficient of the saccade number to fixation number which is outside of the 0.5–3.5 range. The values were assumed arbitrarily and they are derived based on eye tracking recording practice.

The procedure of the reading experiment was basically similar as that of Perceptiveness and Speed Tests (PTs).

The following variants of acoustic conditions were simulated in laboratory tests:

W1 - without the presentation of acoustic stimuli
W2 - with acoustic stimuli - sounds coming from office equipment
W3 - with acoustic stimuli - sounds from office equipment with a quiet conversation in the background (the Speech Transmission Index STI < 0.3)
W4 - with acoustic stimuli - sounds from office equipment with a loud conversation nearby (the Speech Transmission Index STI > 0.45)
W5 - with acoustic stimuli - filtered pink noise.

In each case, the participants of the study were wearing headphones in which sound stimuli were presented. Variants from W2 to W5 have been developed with the assumption that for each of them the equivalent sound level A is 55 dB. The sound level in the headphones was controlled with the MIRE technique using a 25S microphone probe and a sound level meter.

The time of examining the participant in one variant was about 15 minutes. Participants had 15-minute rest breaks between tests. The order of study variants was random - different for each of the subjects.

### Analysis of data

Eye-tracking parameters was registered and analysed for Perceptiveness and Speed Tests (PTs) and for reading the text of a fictional biography, and then to answer questions about the reading (Table 1).

**Table 1.**
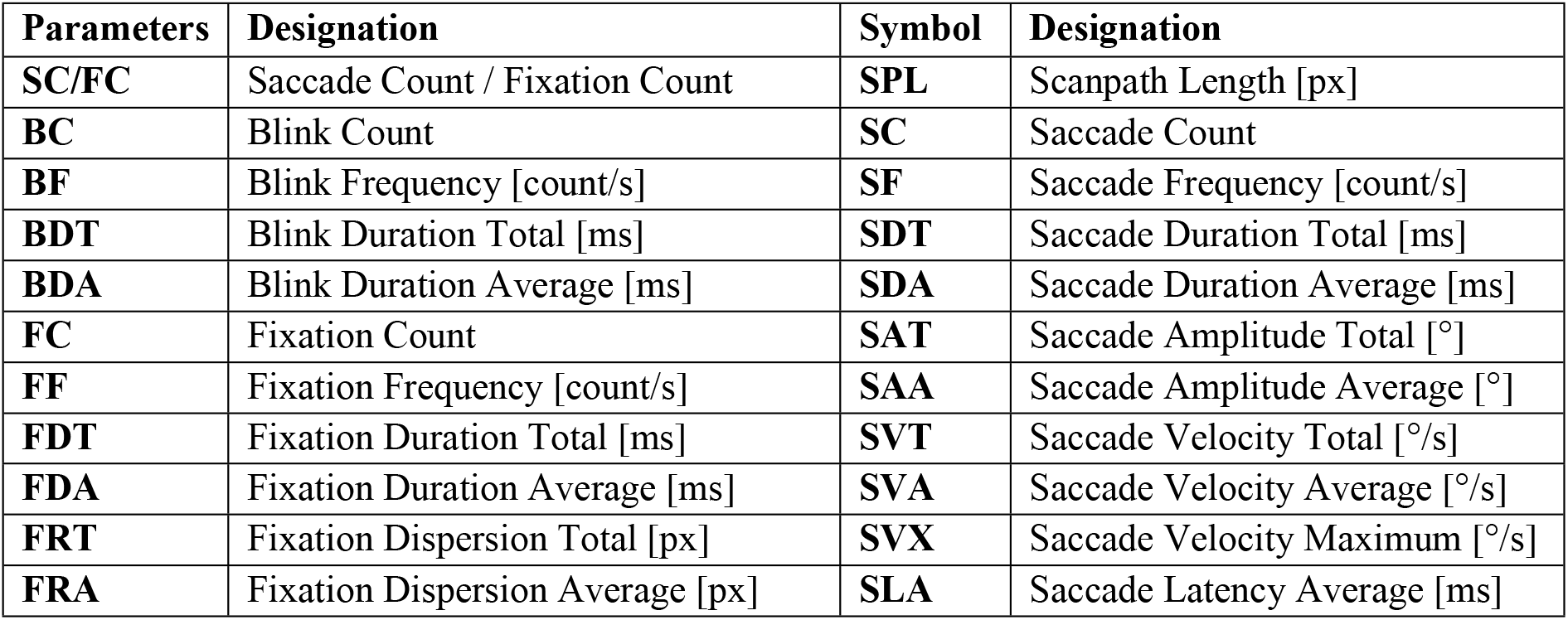
Analysed eye movement parameters

Larger eye defects may cause the loss of some data, therefore, prior to the statistical analysis of parameters, initial numerical data processing is carried out, aimed at, among others, removing artefacts or removing test results for which two assumptions have not been met: Tracking Ratio <90% and the value of the coefficient the number of saccades to the number of fixations not in the range from 0.5 to 3.5. These values were adopted arbitrarily, they result mainly from the practice of eye-tracking registrations and comments contained in reviews of research works in the field of eye-tracking.

Efficiency is a concept that is not clearly defined. In the sciences of organization and management, effectiveness is associated with such concepts as efficiency, efficiency, productivity, profitability, efficiency and even rationality. The purposive approach treats effectiveness as an activity carried out to achieve a specific goal. The systems approach focuses on the relationship between output and input. To assess the tasks performed by the subjects in this study, the following parameters, listed below, were adopted, which are related to the efficiency of performing tasks during mental work. To assess the effectiveness of work in the text: EF-ER-BIO - number of incorrect answers (for questions to the text), EF-DN-BIO - number of “I don’t know” answers (sum of questions P1-P3), EF-IC-BIO - total number of wrong answers and “I don’t know” answers (sum of questions P1-P3) and for tests Numbers, Letters, Alpha, Fractions: EF-ER – error coefficient calculated from the formula: EF- ER=|100-(number of characters counted by the tested person)/(number of all characters to be found in the test)*100|.

The normality of the distribution was analysed by the W Shapiro-Wilk test. The analysis of differences between the individual test variants was carried out on the basis of the analysis of variance with Friedman ANOVA test. Homogeneity of variance was analysed using Levene’s test.

## Results

### Subjective acoustic condition assessment

After each variant, the subjects answered a question about the subjective feeling of the acoustic conditions. On this basis, it was assessed that the individual variants are not too burdensome for the test subjects, but the degree of feeling the nuisance of the acoustic conditions varies depending on the test variant (Table 2.).

**Table 2.** Results of the post-hoc test the Speech Transmission Index)

|  | **Variants of acoustic conditions** |  |  |  |  |  |  |  |  |  |
| --- | --- | --- | --- | --- | --- | --- | --- | --- | --- | --- |
| **Variable** | W1-W2 | W1-W3 | W1-W4 | W1-W5 | W2-W3 | W2-W4 | W2-W5 | W3-W4 | W3-W5 | W4-W5 |
| **ACC** | <0,001** | <0,001** | <0,001** | <0,001** | 0,945 | 0,041* | 0,453 | 0,035* | 0,496 | 0,006** |

\* means $p < 0.05$ ;
\*\* means $p < 0.01$

Conditions in option W4 were assessed as the most onerous (Fig. 2) - only for this variant the level of onerousness exceeded half of the scale. The conditions in variants W2, W3 and W4 were set at a similar level of nuisance. These three options are statistically significantly more onerous than option W1 and statistically significantly less onerous than option W4. The conditions in variant W1, which are significantly better than all other test variants, were considered the best.

The respondents assessed that the conditions of mental work during which the sounds of office equipment are heard with loud conversation are the most onerous, and work in silence was considered the best.

### Visual perception results

Visual perception assessed by eye-tracking parameters while reading the text in different acoustic conditions do not differ statistically significantly (Table 3.).

**Table 3.** Differences in analysed eye-tracking parameters

| Variables | Text reading |  | Sub-test of Digits |  | Sub-test of Letters |  | Alfa Sub-test |  | Sub-test of Fractions |  |
| --- | --- | --- | --- | --- | --- | --- | --- | --- | --- | --- |
|  | F | p | F | p | F | p | F | p | F | p |
| SC/FC | 0,39 | 0,815 | 0,17 | 0,951 | 8,40 | 0,078 | 0,71 | 0,586 | 0,20 | 0,938 |
| BC | 0,25 | 0,908 | 0,12 | 0,975 | 4,80 | 0,308 | 0,16 | 0,956 | 0,17 | 0,955 |
| BF | 0,27 | 0,898 | 0,75 | 0,558 | 0,40 | 0,807 | 0,82 | 0,512 | 0,12 | 0,976 |
| BDT | 0,51 | 0,730 | 0,40 | 0,810 | 0,07 | 0,992 | 0,35 | 0,845 | 0,11 | 0,980 |
| BDA | 0,32 | 0,867 | 3,09 | **0,021\*** | 0,54 | 0,707 | 0,28 | 0,891 | 2,65 | **0,040\*** |
| FC | 0,43 | 0,784 | 0,14 | 0,968 | 0,36 | 0,835 | 0,25 | 0,912 | 2,28 | 0,063 |
| FF | 0,14 | 0,967 | 0,15 | 0,963 | 0,49 | 0,740 | 0,35 | 0,845 | 1,95 | 0,104 |
| FDT | 0,46 | 0,766 | 0,25 | 0,911 | 0,97 | 0,425 | 1,19 | 0,318 | 0,55 | 0,702 |
| FDA | 0,26 | 0,906 | 0,14 | 0,967 | 0,50 | 0,733 | 0,11 | 0,981 | 2,70 | **0,032\*** |
| FRT | 0,04 | 0,996 | 0,29 | 0,884 | 0,22 | 0,925 | 7,90 | 0,095 | 0,14 | 0,969 |
| FRA | 0,24 | 0,913 | 0,16 | 0,957 | 0,21 | 0,934 | 1,52 | 0,198 | 0,62 | 0,650 |
| SPL | 0,22 | 0,929 | 0,67 | 0,613 | 1,24 | 0,296 | 0,67 | 0,612 | 0,59 | 0,672 |
| SC | 0,47 | 0,759 | 0,55 | 0,702 | 0,28 | 0,888 | 0,39 | 0,818 | 1,33 | 0,261 |
| SF | 0,61 | 0,653 | 0,53 | 0,712 | 0,27 | 0,900 | 0,34 | 0,849 | 12,91 | **0,012\*** |
| SDT | 0,50 | 0,739 | 0,02 | 0,999 | 1,21 | 0,308 | 0,64 | 0,637 | 0,32 | 0,866 |
| SDA | 0,53 | 0,714 | 0,04 | 0,996 | 0,25 | 0,910 | 0,35 | 0,844 | 2,04 | 0,728 |
| SAT | 0,08 | 0,989 | 0,32 | 0,866 | 0,17 | 0,953 | 0,77 | 0,549 | 0,32 | 0,866 |
| SAA | 0,75 | 0,558 | 1,08 | 0,370 | 0,09 | 0,985 | 1,05 | 0,385 | 0,27 | 0,896 |
| SVT | 0,33 | 0,860 | 3,22 | 0,521 | 1,02 | 0,399 | 0,30 | 0,877 | 0,97 | 0,427 |
| SVA | 0,89 | 0,471 | 0,66 | 0,620 | 0,84 | 0,504 | 0,50 | 0,738 | 0,09 | 0,985 |
| SVX | 1,17 | 0,324 | 0,08 | 0,990 | 5,11 | 0,276 | 0,44 | 0,777 | 1,93 | 0,108 |
| SLA | 0,35 | 0,847 | 0,12 | 0,977 | 0,57 | 0,682 | 0,54 | 0,710 | 1,12 | 0,348 |

\* means $p < 0.05$

In the PTs test, the occurrence of statistically significant differences between the variants was found in the Sub-test of Digits (parameter BDA) and the Sub-test of Fractions (for BDA, FDA and SF parameters).

The occurrence of statistically significant differences between the variants was found: in Sub-test of Digits for the BDA and parameter Sub-test of Fractions for BDA, FDA and SF parameters (Table 4.). Below are graphs (Fig 4 - 5) for parameters and tests for which statistical analysis showed significant differences between the variants. With regard to blinking, there are differences in the Digits and Fractions test, but it is difficult to clearly determine the trends. The occurring differences are rather accidental.

**Table 4.** Results of the post-hoc test of variables for which statistically significant results of the analysis of variance were obtained

|  | Variables | Variants of acoustic conditions |  |  |  |  |  |  |  |  |  |
| --- | --- | --- | --- | --- | --- | --- | --- | --- | --- | --- | --- |
|  |  | W1-W2 | W1-W3 | W1-W4 | W1-W5 | W2-W3 | W2-W4 | W2-W5 | W3-W4 | W3-W5 | W4-W5 |
| Sub-test<br>of Digits | BDA | 0,658 | **0,018\*** | 0,385 | 0,921 | **0,007\*\*** | 0,674 | 0,572 | **0,002\*** | **0,017\*** | 0,311 |
| Sub-test<br>of<br>Fractions | BDA | 0,499 | 0,171 | **0,006\*\*** | 0,985 | 0,503 | 0,041 | 0,506 | 0,161 | 0,184 | **0,008\*\*** |
|  | FDA | 0,697 | **0,017\*** | 0,450 | 0,085 | **0,046\*** | 0,713 | 0,185 | 0,105 | 0,486 | 0,343 |
|  | SF | 0,409 | 0,227 | 0,364 | 0,773 | 0,636 | 0,546 | **0,042\*** | 0,591 | 0,546 | **0,036\*** |

\* means $p < 0.05$ ; \*\* means $p < 0.01$

**Fig. 4.**
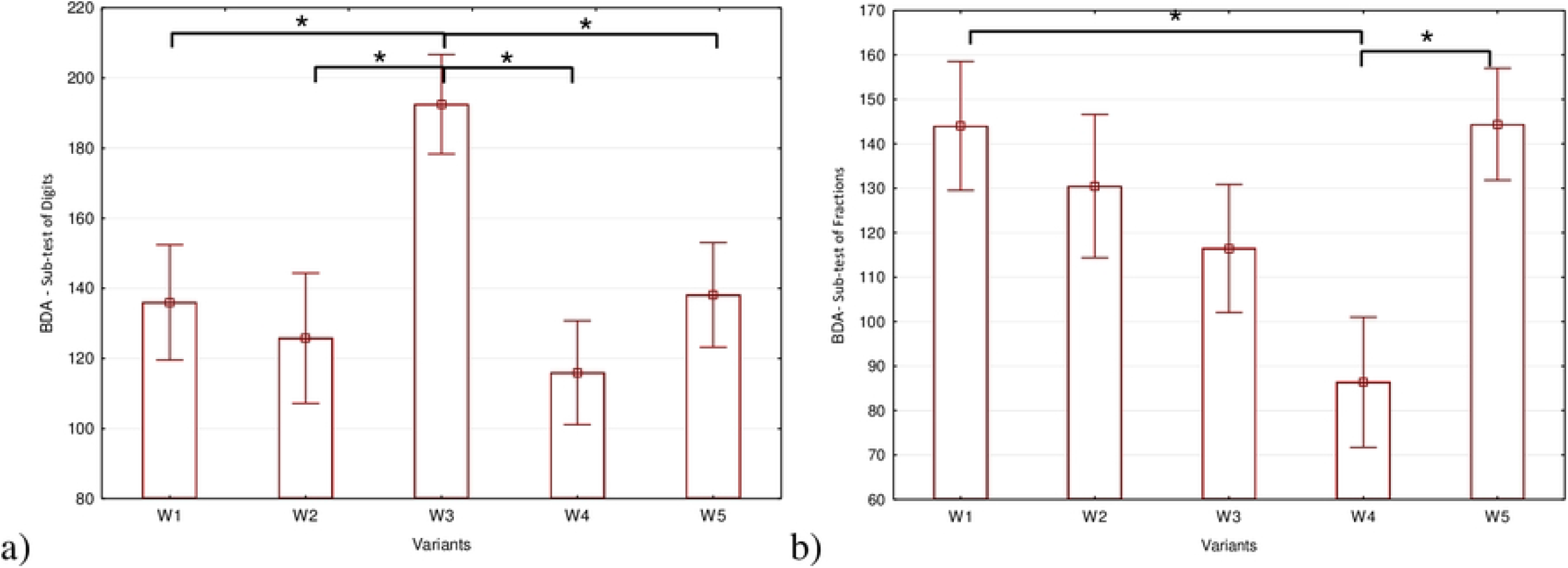
The mean values and errors of parameter BDA (Blink Duration Average) in Sub-test of Digits (a) and Sub-test of Fractions (b) caused by acoustic conditions.

**Fig. 5.**
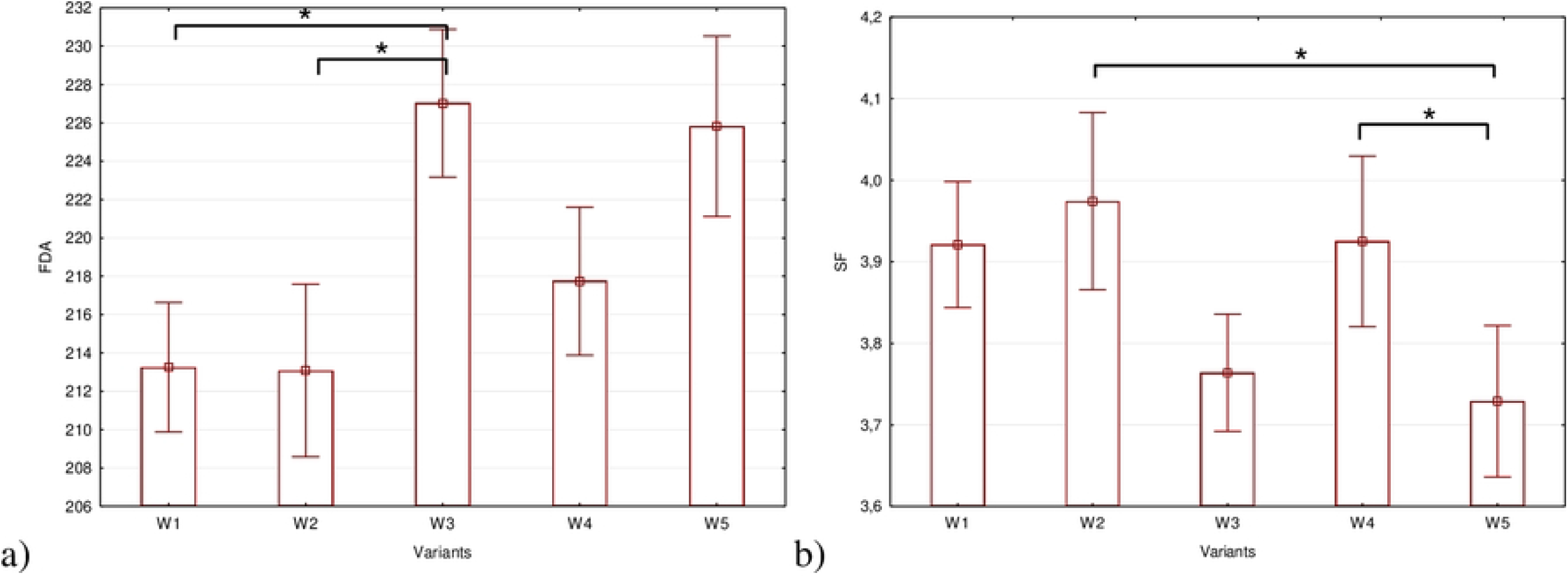
The mean values and errors of parameter FDA (Fixation Duration Average) (a) and SF (Saccade Frequency) (b) in Sub-test of Fractions caused by acoustic conditions.

### Workload

The workload assessed by NASA-TLX scales did not differ between variants, no statistically significant differences were found. (Table 5.).

**Table 5.** Differences in analysed NASA-TLX scales

| NASA-TLX scales | F | p |
| --- | --- | --- |
| **Mental Demand** | 1,12 | 0,3474 |
| **Physical Demand** | 0,22 | 0,9289 |
| **Temporal Demand** | 0,18 | 0,9460 |
| **Performance** | 0,90 | 0,4672 |
| **Effort** | 0,26 | 0,9022 |
| **Frustration** | 0,42 | 0,7944 |

\* means $p < 0.05$ ; \*\* means $p < 0.01$

The assessment using the NASA-TLX questionnaire indicates that the subjects similarly assessed the level of mental load, physical load, time pressure, efficiency, effort and frustration, regardless of the acoustic conditions in which mental work was performed.

### Work efficiency results

During reading the text, there were statistically significant differences in efficiency results (the EF-IC-BIO parameter) between variant W1 and variant W4 and between variant W3 and W4 (Table 6.). The number of incorrect answers and “I don’t know” combined was statistically significantly higher in the variant with audible sounds of office equipment with loud conversation (W4) than in the variant with audible sounds of office equipment with quiet conversation (W3) and in the variant without the presentation of acoustic stimuli (W1) – Table 7, Fig 6.

**Table 6.** Differences in analysed Work efficiency results

|  | **Parameter** | **F** | **p** |
| --- | --- | --- | --- |
| **Text reading** | EF-ER-BIO | 2,40 | 0,0519 |
|  | EF-DN-BIO | 18,21 | 0,0011* |
|  | EF-IC-BIO | 3,42 | 0,0100* |
| **Sub-test of Digits** | EF-ER | 0,54 | 0,7051 |
|  | EF-CR | 9,13 | 0,0579 |
| **Sub-test of Letters** | EF-ER | 0,53 | 0,7110 |
|  | EF-CR | 3,69 | 0,4507 |
| **Alfa Sub-test** | EF-ER | 0,45 | 0,7756 |
|  | EF-CR | 5,77 | 0,2170 |
| **Sub-test of Fractions** | EF-ER | 32,29 | 0,0000** |
|  | EF-CR | 20,19 | 0,0005** |

\* means $p < 0.05$ ; \*\* means $p < 0.01$

**Table 7.** Results of the post-hoc test of variables for which statistically significant results of the analysis of variance were obtained

| Test | Variable | Variants of acoustic conditions |  |  |  |  |  |  |  |  |  |
| --- | --- | --- | --- | --- | --- | --- | --- | --- | --- | --- | --- |
|  |  | W1-W2 | W1-W3 | W1-W4 | W1-W5 | W2-W3 | W2-W4 | W2-W5 | W3-W4 | W3-W5 | W4-W5 |
| Text reading | EF-DN-BIO | 0,1538 | 0,4744 | 0,3333 | 0,4872 | 0,3205 | 0,4872 | 0,6410 | 0,8077 | 0,9615 | 0,1538 |
|  | EF-IC-BIO | 0,1463 | 0,7157 | **0,0040\*** | 0,1827 | 0,0698 | 0,1463 | 0,9034 | **0,0012\*** | 0,0904 | 0,1157 |
| Sub-test of Fractions | EF-ER | 0,4257 | 0,8051 | **<0,001\*\*** | **<0,001\*\*** | 0,2972 | **<0,001\*\*** | **<0,001\*\*** | **<0,001\*\*** | **<0,001\*\*** | 0,2918 |
|  | EF-CR | 0,3205 | 0,1923 | 0,6410 | 0,7051 | 0,1282 | 0,3205 | 0,3846 | 0,4487 | 0,5128 | 0,0641 |

\* means $p < 0.05$ ; \*\* means $p < 0.01$

**Fig. 6.**
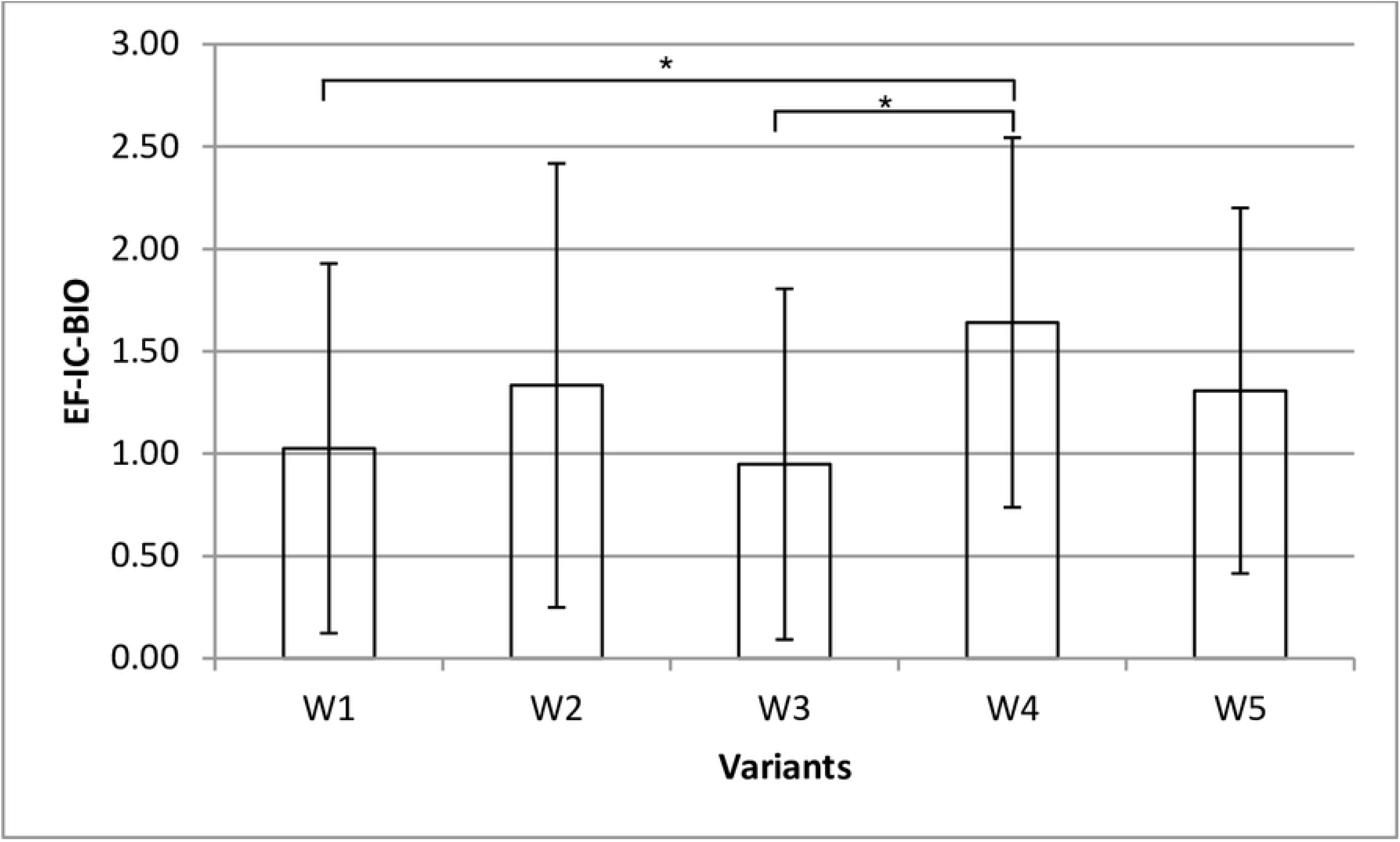
The mean values and errors of parameter EF-IC-BIO - total number of wrong answers and “I don’t know” answers (sum from all questions) in acoustic conditions.

Of the 4 TWAP tests, only in the Fractions Sub-test (the most difficult one, because during the search the test person has to check both the numerator and the denominator of a fraction) statistically significant differences were found (Tab 5 and Tab 6.). Most errors were made during mental work with the audible sounds of office equipment with a loud conversation (Variant W4) and during mental work performed during audible filtered pink noise (Variant W5). The number of errors made in these variants was about twice as high as in the other test variants (Fig. 7.). The results of work efficiency in the Digits, Letters, Alpha tests did not differ depending on the acoustic conditions.

**Fig. 7.**
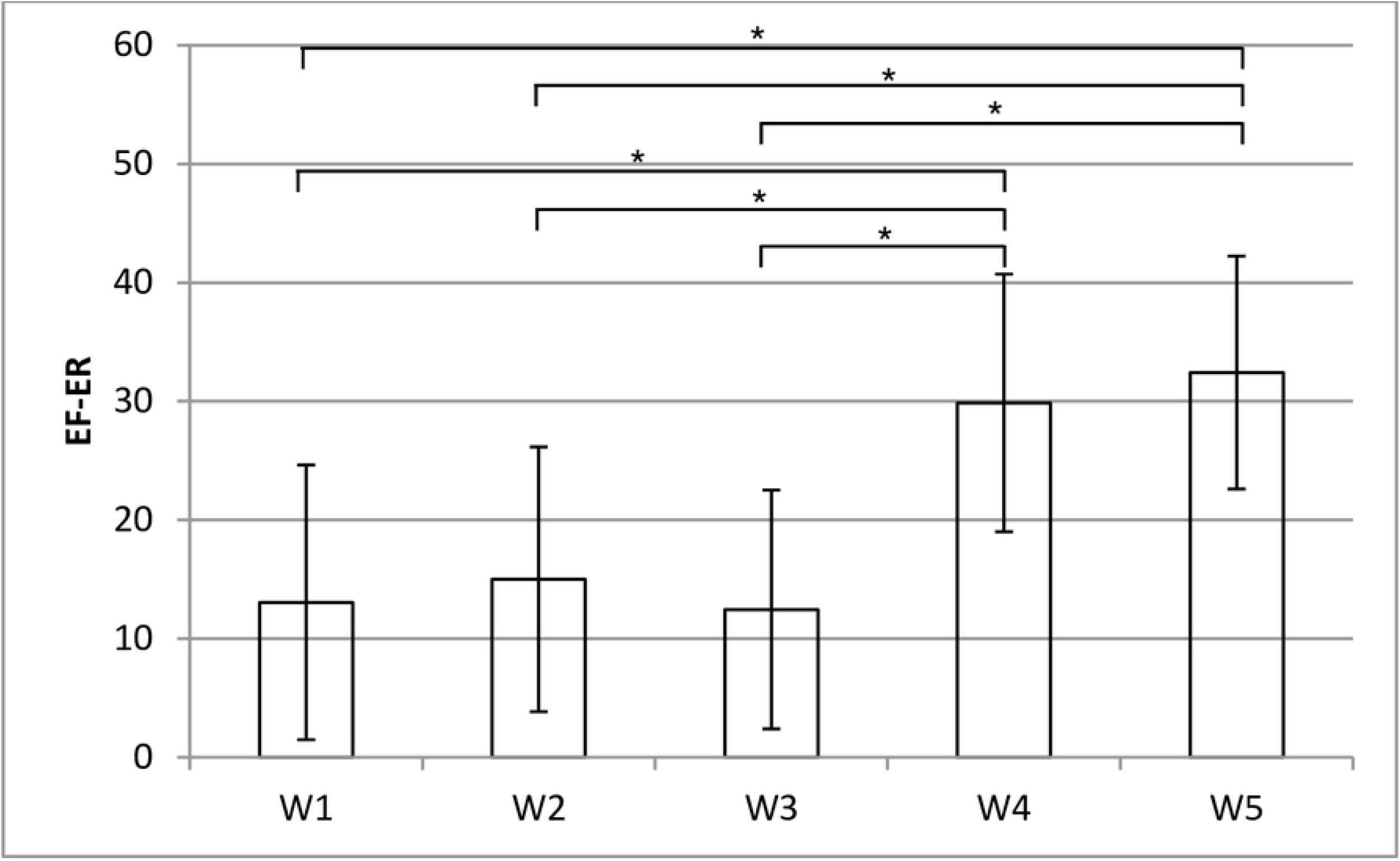
The mean values and errors of parameter EF-ER in Sub-test of Fractions – error coefficient in acoustic conditions.

## Discussion and conclusions

The present study investigated the impact of the type of acoustic conditions, typical of office work, on visual perception (eye-tracking parameters), in comparison to subjectively assessed workload, subjective acoustic condition assessment and work efficiency.

The subjective assessment of the acoustic conditions nuisance (ACC) carried out after each test variant showed that there are significant differences in the perception of individual test variants. The respondents assessed that the conditions of mental work during which the sounds of office equipment are heard with loud conversation are the most onerous (Variant W4), and the work in Variant W1 - without the presentation of acoustic stimuli - was considered the best. Variant W4 (with sounds from office equipment with a loud conversation nearby (STI > 0.45) was rated as more cumbersome compared to all other variants. Statistically significant differences occurred between variants W3 and W4, which had the same equivalent sound level A (55 dB) and differed only in the value of STI. Higher workload was observed under conditions of higher speech intelligibility. Also in other studies [28,29] a higher workload under conditions of maximum speech intelligibility was observed.

At the same time, no influence of acoustic conditions on the subjective assessment of the workload was observed. The assessment using the NASA-TLX questionnaire indicates that participants similarly assessed the level of mental load, physical load, time pressure, efficiency, effort, and frustration, regardless of the acoustic conditions in which mental work was performed. Also, the study [30] found that noise-related dissatisfaction was correlated with dissatisfaction with the environment and work, although no relationship was found with job performance. In our measurements, inter-individual variability of the workload appeared. Research conducted by Ebissou et al. [28] indicates that some people may be insensitive to speech intelligibility because of the strong inter-individual variability. This may explain the lack of statistically significant differences between variants in our research. As shown in Fig. 8 cumulative TLX scores achieved by different individuals (p1-p39) varied greatly.

**Fig. 8.**
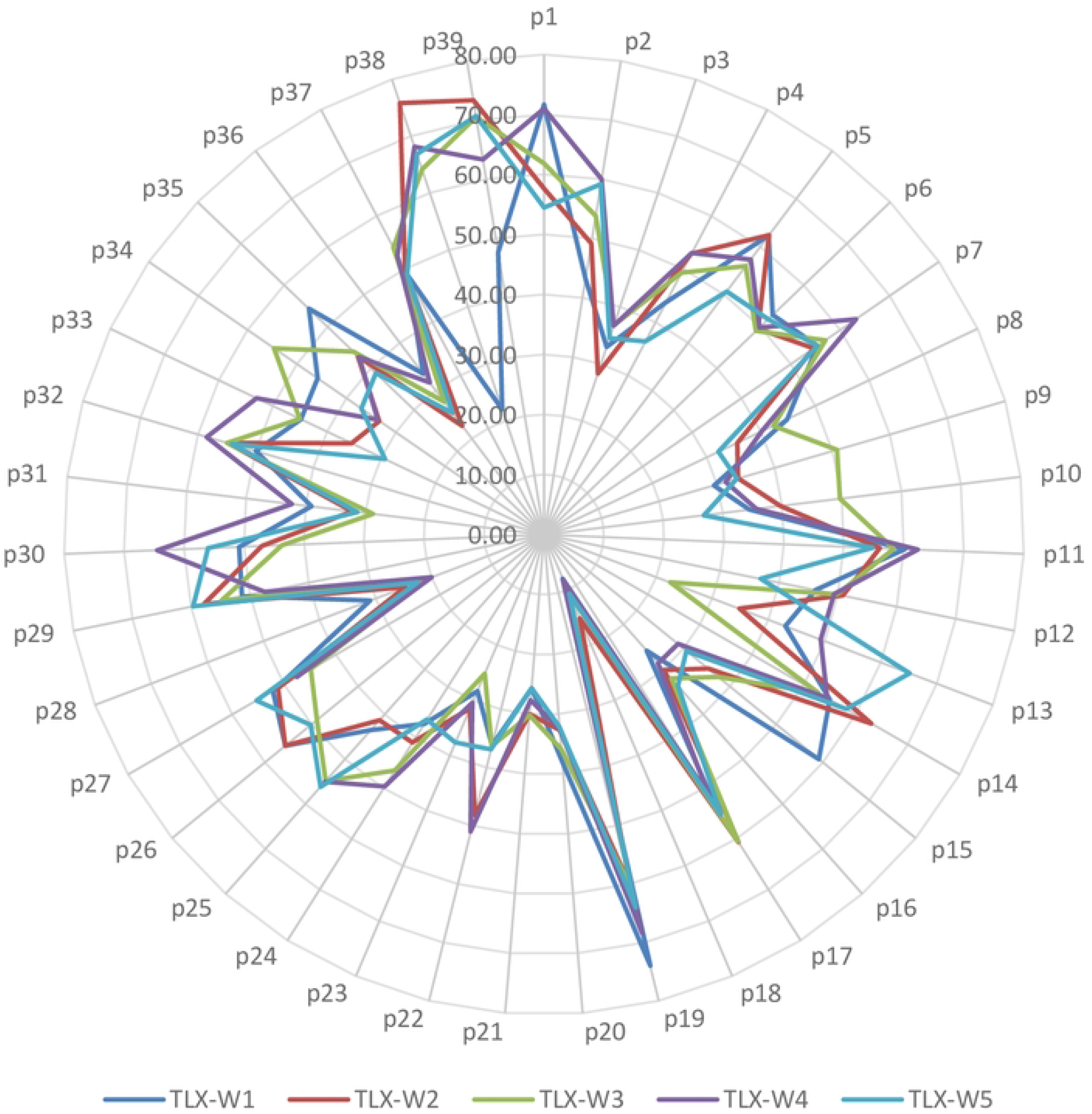
The results of the workload assessed using the NASA-TLX questionnaire in various acoustic conditions (W1-W5) achieved by the subjects (p1-p39)

The assessment of work efficiency in the reading test indicates the existence of statistically significant differences. The number of incorrect answers and “don’t know” answers combined (parameter EF-IC-BIO) was statistically significantly higher in the variant with audible sounds of office equipment with loud conversation (W4) than in the variant with audible sounds of office equipment with quiet conversation (W3) and in the variant without the presentation of acoustic stimuli (W1). Research [13] also indicate an increase in the number of errors when performing tasks with a higher level of difficulty. Of the 4 PTs tests, only in the Fractions test (the most difficult one, because during the search the test person has to check both the numerator and the denominator of a fraction) statistically significant differences were found. Most errors were made during mental work with the audible sounds of office equipment with a loud conversation (Variant W4) and during mental work performed during audible filtered pink noise (Variant W5). The number of errors made in these variants was about twice as high as in the other test variants. The results of work efficiency in the Digits, Letters and Alpha tests did not differ depending on the acoustic conditions. Such results are also consistent with research [13], which proved that when the tasks were medium or difficult, their performance was more strongly influenced by the type of noise, not its level. In our experiment, the W2-W5 variants did not differ in sound level (the equivalent sound level A was 55 dB) but in its type.

Oculomotor parameters while reading the text in different acoustic conditions do not differ statistically significantly. In turn, in the PTs test, the occurrence of statistically significant differences between the variants was found in the Digit test (for parameters related to blinking: BDA, BDX, BDM) and the Fraction test (for BDA, FDA and SF parameters). As proved in the publication [31] perceptual load and cognitive load had an opposite effect on fixation duration. Higher perceptual load led to decreased fixation durations but higher cognitive load led to increased fixation durations. During the experiment, the subjects had to perform the same task in different acoustic conditions. A longer fixation duration in the Fraction test indicates that variant W3 caused a greater mental load than in variants W1 and W2. At the same time, the lack of statistically significant differences in other tests and when reading the text, indicates that the acoustic conditions do not affect visual perception.

The differences between the test variants in the value of the BDA parameter are not unambiguous. In the Sub-test of Digits, the longest blinking times were observed in the W3 variant, and in the Sub-test of Fractions - in the W1 and W5 variants. In research greater blink duration or eye blink rate was indicated as a parameter of mental workload and the eye blink rate can be an indicator of effective operation under high incentive conditions. [32,33,34]. However, the actual direction of this relation is uncertain.

In conclusion, cognitive performance depends on the type of noise. Acoustic factors had an interactive effect on workers cognitive functions. Participants also experienced higher levels of discomfort in conditions of higher speech intelligibility. Research also indicates an increase in the number of errors when performing tasks with a higher level of difficulty. The most errors were made during mental work with the audible sounds of office equipment with a loud conversation and during mental work performed with audible filtered pink noise.

The lack of statistically significant differences in almost all perceptiveness tests (except the most difficult one) and during text reading indicates that the acoustic conditions do not affect visual perception but rather cognitive performance. More studies are needed to further explore this relationship.

